# Reproducible functional connectivity endophenotype confers high risk of ASD diagnosis in a subset of individuals

**DOI:** 10.1101/2020.06.01.127688

**Authors:** Sebastian GW. Urchs, Hien Duy Nguyen, Clara Moreau, Christian Dansereau, Angela Tam, Alan C. Evans, Pierre Bellec

**Affiliations:** Montreal Neurological Institute and Hospital, McGill University, 3801 Rue de l’Université, QC H3A 2B4, Montreal, Canada; Centre de Recherche de l’Institut Universitaire de Gériatrie de Montréal, 4565 Queen Mary Rd, QC H3W 1W5, Montreal, Canada; Department of Mathematics and Statistics, La Trobe University, Plenty Rd & Kingsbury Dr, VIC 3086, Bundoora, Australia, Centre de Recherche de l’Institut Universitaire en Santé Mentale de Montréal, 7401 Rue Hochelaga, QC H1N 3M5, Montreal, Canada; Sainte Justine Research Center, University of Montreal, 3175 Chemin de la Côte-Sainte-Catherine, QC H3T 1C5, Montreal, Canada

## Abstract

Functional connectivity (FC) analyses of individuals with autism spectrum disorder (ASD) have established robust alterations of brain connectivity at the group level. Yet, the translation of these imaging findings into robust markers of individual risk is hampered by the extensive heterogeneity among ASD individuals. Here, we report an FC endophenotype that confers a greater than 7-fold risk increase of ASD diagnosis, yet is still identified in an estimated 1 in 200 individuals in the general population. By focusing on a subset of individuals with ASD and highly predictive FC alterations, we achieved a greater than 3-fold increase in risk over previous predictive models. The identified FC risk endophenotype was characterized by underconnectivity of transmodal brain networks and generalized to independent data. Our results demonstrate the ability of a highly targeted prediction model to meaningfully decompose part of the heterogeneity of the autism spectrum. The identified FC signature may help better delineate the multitude of etiological pathways and behavioural symptoms that challenge our understanding of the autism spectrum.

## Introduction

### Background

Autism spectrum disorder (ASD) is a neurodevelopmental disorder diagnosed in more than 1% of children (***Bai et al., 2019***) that is defined by impairments of social interaction and repetitive behaviour (***American Psychiatric Association. and DSM-5 Task Force., 2013***), and has been linked to alterations of brain organization (***Holiga et al., 2019***) and genetics (***Grove et al., 2019***). A core goal of clinical neuroscience is to understand the neurobiological etiology of this complex and heterogeneous disorder (***Lombardo et al., 2019***) by identifying reliable neurobiological endophenotypes that confer a high risk of the disorder and are sufficiently common to be investigated in large cohort studies. To date, existing genetic risk markers of ASD are either extremely rare (e.g. monogenic disorders, ***de la Torre-Ubieta et al., 2016***) or convey only a very low individual risk of the disorder (e.g. common genetic risk factors, ***Sanders et al., 2015***). Current neuroimaging based efforts to identify brain based endophenotypes that can predict ASD (***Abraham et al., 2017***; ***Heinsfeld et al., 2018***) have limited accuracy, likely due to the extensive heterogeneity of the disorder (***Lombardo et al., 2019***; ***Jacob et al., 2019***, see also ***Figure 2***). Although it may not currently be possible to identify a single brain based risk marker that is highly predictive of an ASD diagnosis in any autistic individual, we may be able to do so for a subset of autistic individuals. Transductive conformal prediction (TCP, ***Vapnik, 1998***; ***Vovk et al., 2005***) is a promising statistical framework for this purpose, that has been successfully applied to predict the clinical status of depressed patients from neuroimaging data, previously (***Nouretdinov et al., 2011***). TCP explicitly computes the confidence with which a clinical label (i.e. ASD or neurotypical control, NTC) can be predicted for each individual, and we can use this estimate to limit our model to predict only those individuals for whom we have very high confidence of their ASD diagnosis. Our main goal is to apply TCP to identify resting-state functional Magnetic Resonance Imaging (fMRI) based ASD-endophenotypes associated with high risk and relatively high prevalence.

**Figure 1.**
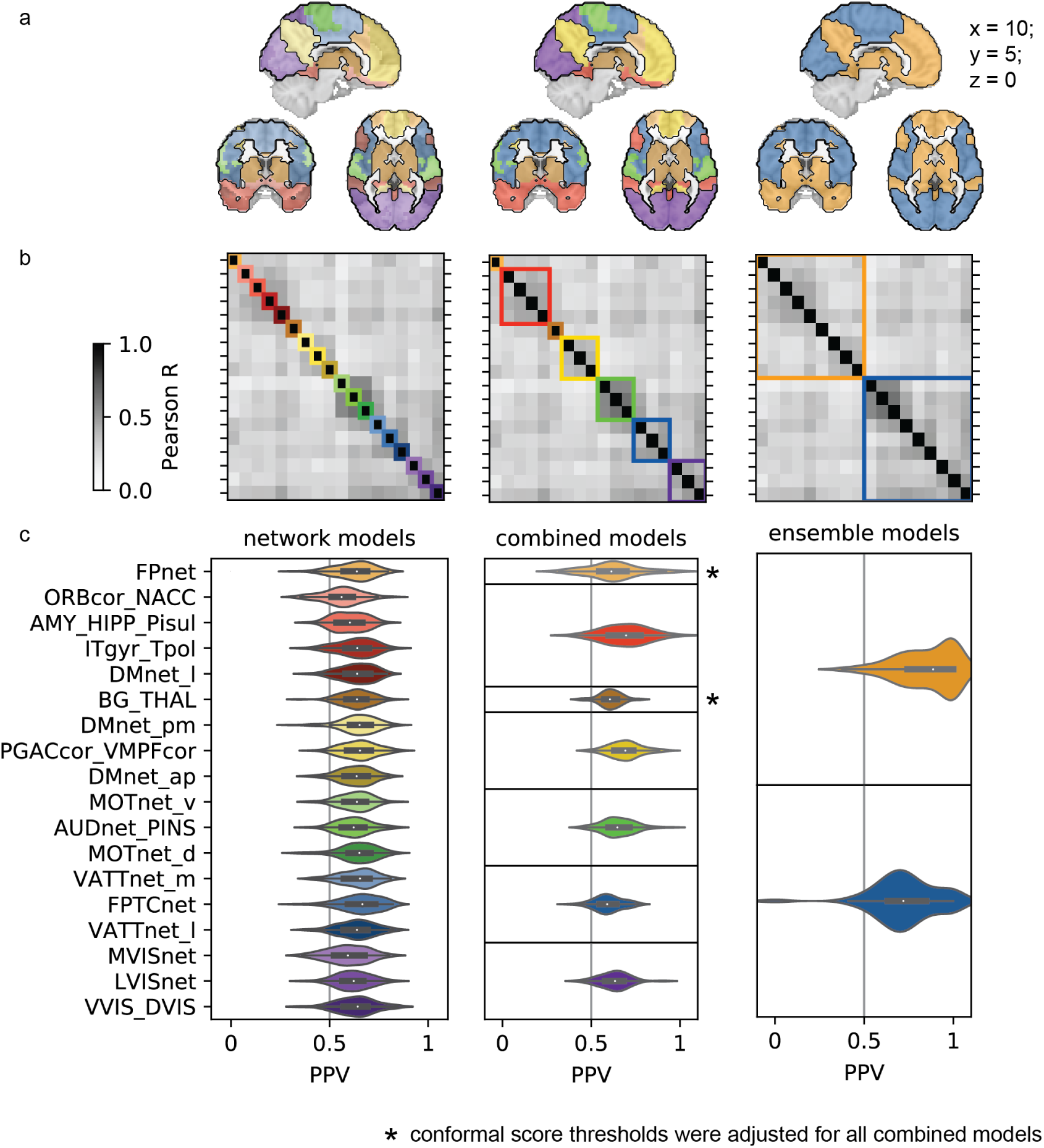
Combining network predictors with correlated conformal scores results in higher prediction performance. Conformal predictions based on individual networks (left column) were not associated with high PPV (c). The predicted ASD conformal scores were correlated between networks (b) and were used to cluster networks into 7 combined predictors (middle column). Clusters predominantly broke down along boundaries of large scale functional brain networks (a). Networks with correlated conformal predictions were further clustered into two large ensemble predictors (right column), that combined predominantly unimodal (blue) and transmodal (orange) brain networks respectively (a, right column). Predictions of the ensemble of more transmodal networks (orange) gave rise to a high risk signature (HRS) that predicted ASD with high PPV (c, top).

**Figure 2.**
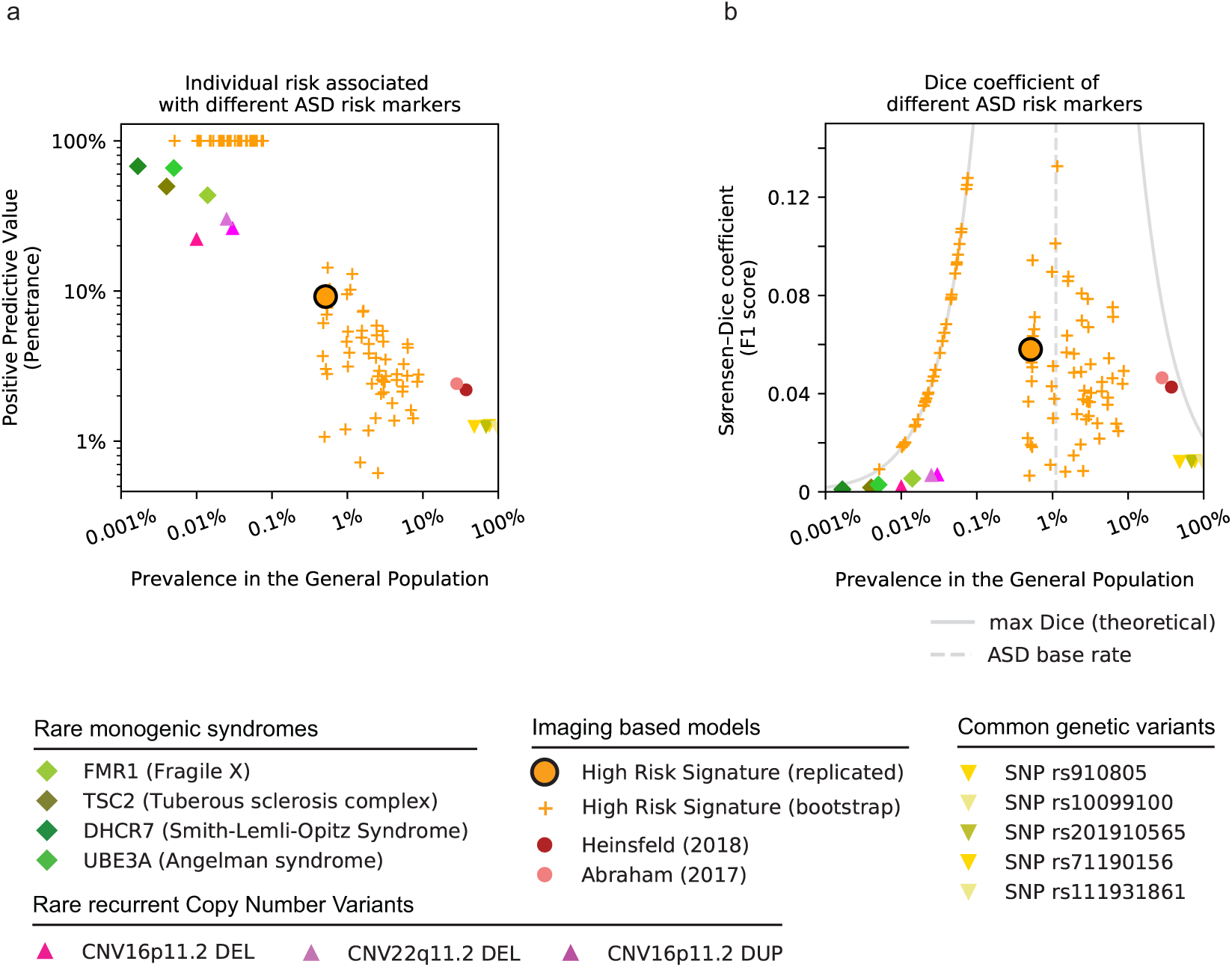
HRS is more common than genetic risk markers and confers higher risk than traditional imaging models. a) Monogenic syndromes (green rhombs) and recurrent Copy Number Variants (pink triangles) confer high risk of ASD diagnosis (vertical axis), but are rare (horizontal axis). ASD related single nucleotide polymorphisms (yellow triangles) are very common, but confer negligible risk of ASD. Current imaging based predictive models (pink circles) identify large portions of the general population with low risk of ASD. The high risk ASD signature (orange, black outline) identifies a small portion of the general population with elevated risk of ASD diagnosis, concordant with the estimated performance in the discovery data (orange plus signs). b) The Dice coefficient (***Equation 9***) reflects the degree of overlap between individuals identified by a risk marker and the true ASD population. Rare genetic risk factors (green rhombs and pink triangles) with high PPV identify a small subset of ASD individuals and thus have low Dice coefficients. Common ASD related genetic variants (yellow triangles) identify large portions of the general population but with low PPV and thus also have low Dice coefficients. The HRS shows Dice coefficients that are comparable but higher than those of existing imaging models, reflecting the large increase in PPV over these models and the lower sensitivity.

### Genetic risk markers of ASD

ASD is a highly heritable disorder and to date, the best established ASD risk factors are genetic markers. A recent multi-national study of more than 2 million individuals estimated the heritability of ASD at 80% (***Bai et al., 2019***). Both common (e.g. Single-nucleotide polymorphism, SNP, ***Grove et al., 2019***) and rare genetic variants (e.g. recurrent Copy-number variant, CNV, ***Sanders et al., 2019***) have been shown to contribute to the genetic etiology of ASD (***Geschwind and State, 2015***). Several monogenic syndromes have also been associated with a very high risk of autism-like symptoms (i.e. in more than 30% of individuals with the syndrome), but these disorders are exceedingly rare in the general population, typically detected in fewer than 0.01% of individuals (***de la Torre-Ubieta et al., 2016***). By comparison, ASD is diagnosed relatively frequently in about 1% of individuals in the general population (***Bai et al., 2019***). Only five common genetic variants (found in more than 5% of the general population) have recently been robustly associated with ASD through genome-wide association studies (***Grove et al., 2019***). However, each of these common variants increase the odds of an ASD diagnosis only minimally in carriers compared to non-carriers (i.e. the Odds Ratio is approximately 1.2 or close to equal, see ***Equation 8***). Nevertheless, common genetic variants are thought to account for a large part of genetic ASD liability, with estimates ranging between 20% (***Robinson et al., 2016***) and 50% (***Gaugler et al., 2014***). In between the rare, high risk monogenic disorders and the common, but low risk genetic variants, sits a gap of knowledge that has been labeled the “missing heritability” (***Manolio et al., 2009***; ***Maher, 2008***). The very large sample sizes necessary (***Khera et al., 2018***) to robustly identify the likely polygenic interaction effects (***O’Connor et al., 2019***) pose a challenging limitation that makes the identification of common, high risk genetic factors of ASD difficult.

### Neuroimaging based risk markers of ASD

Functional magnetic resonance imaging (fMRI) measures the functional connectivity (FC) between brain regions and has been shown to be sensitive to changes in the functional brain organization in ASD (***Castellanos et al., 2013***; ***Holiga et al., 2019***). Recent work has therefore used high-dimensional FC measures to predict the clinical ASD diagnosis of individuals (***Abraham et al., 2017***; ***Heinsfeld et al., 2018***; ***Yahata et al., 2016***). These models make a prediction for every individual in a data set and seek to optimize the accuracy of all predictions. That is, they give equal importance to correctly identifying an individual with ASD (sensitivity, see ***Equation 1***) and to correctly not identifying a NTC individual (specificity, see ***Equation 2***). As a consequence, predictions by these models typically have balanced sensitivity and specificity. When such a model is applied to an unselected general population sample, where only very few individuals will truly have ASD (i.e. 1 — 2%), the ability of the model to correctly identify unaffected individuals as not having the condition (specificity) becomes more important. For example, if 20 individuals in a sample of 1000 have ASD (i.e. 980 are healthy), then a model with 70% sensitivity and 70% specificity will correctly identify 14 ASD individuals (20 x sensitivity of 0.7) and correctly not identify 686 healthy individuals (980 x specificity of 0.7). The model will however also incorrectly identify 294 healthy individuals as ASD patients (980 ∗ (1 − 0.7)). This means that only 14 out of 308 (or 4%) individuals identified by the model will truly have ASD. This value is also called the positive predictive value (PPV) of the model and depends on the prevalence of the predicted disorder in the sample (see Methods and ***Equation 3***). The PPV thus reflects the risk of the disorder that a prediction by the model confers for an individual. In the above example, the PPV is only twice as large as the baseline risk of someone we know nothing about (i.e. the prevalence of the disorder in the sample). Recent FC based ASD classification models report sensitivity and specificity estimates that translate to low PPVs of 2.4% to 2.2% in the general population (***Abraham et al., 2017***; ***Heinsfeld et al., 2018***).

### Transductive conformal prediction

Performance metrics such as accuracy, sensitivity, and specificity provide us with some measures regarding the confidence that we can place on the quality of the predictions made by a model, on average, across all observed samples from either a testing or training data set. But what we are particularly interested in is the amount of confidence that we can place in the specific clinical label predicted for each individual. This is akin to the difference between the usual confidence and prediction intervals (***Kümmel et al., 2018***). Whereas the model in the above example identifies hundreds of individuals as ASD patients, maybe for some of the individuals, the degree of “confidence” of the label is more limited, based on the idiosyncrasies of the individual. We may decide that we only want to take a closer look at those individuals for whom the model is very confident of their ASD diagnosis. Conformal prediction is a statistical framework to make explicit the level of confidence that an analyst may have regarding the classification of any particular individual (***Vovk et al., 2005***). Given an individual that we want to classify as either neurotypical or ASD, the conformal predictor asks: “how unusual would this individual be, if they were a neurotypical individual?” and “how unusual would it be, if they were an individual with ASD?”. The predictor then answers each of these questions by comparing the individual to known neurotypical individuals and to known individuals with ASD, respectively. In this way, we will compute two “unusualness” scores for the individual, one for each of the two possible label classes. More technical introductory accounts of the conformal prediction logic can be found in articles such as ***Gammerman and Vovk (2007***), and ***Shafer and Vovk (2008***).

### Objectives

Here we aim to identify FC signatures of ASD that are substantially more common than rare monogenic disorders and carry substantially higher individual risk than current imaging based models of ASD. We hypothesize that by limiting predictions to the most confident cases, we will identify subsets of ASD individuals who share very predictive, high risk FC signatures. We further hypothesize that the FC of different brain networks may give rise to distinct high risk FC signatures. Our objectives are thus to:

1. Identify sets of brain networks with FC profiles highly predictive of ASD diagnosis.
2. Evaluate the identified high risk profiles on an independent dataset, and estimate their prevalence and positive predictive value in the general population.
3. Characterize the connectivity and symptom phenotype of the individuals identified by the high risk FC profiles.

## Results

We investigated whether the seed-based FC maps of 18 functional brain networks could be used to predict ASD diagnosis with high PPV in a subset of individuals. To do so, we estimated how conformal the FC of a new, unclassified individual was compared to individuals with a known ASD diagnosis or NTC. These estimates of conformality then allowed us to make a prediction of ASD diagnosis only for those individuals for whom we had very high confidence that their FC was very atypical for NTC (NTC conformal score < 5%) and not very atypical for ASD (ASD conformal score > 5%). We identified the groups of brain networks that gave rise to the predictions with the highest PPV across bootstrap samples of the discovery data and then tested the generalizability of these predictions in an independent replication dataset.

### Individual networks do not predict ASD with high PPV

We first evaluated the PPV of conformal ASD diagnosis predictions made with high confidence, based on the FC of each of the 18 brain networks. To do so, we computed the median PPV of high confidence conformal predictions for each brain network across 100 bootstrap samples (bootstrap PPV) of the discovery data. The bootstrap PPV of high confidence conformal ASD diagnosis predictions ranged from 56% (orbitofrontal network) to 66% (frontoparietal network) and was 63% on average across all networks. That is, among the individuals predicted with high confidence to have an ASD diagnosis, 63% on average did have an ASD diagnosis. As expected, the predictions were made with high specificity (91% on average across all networks) and low sensitivity (16% across all networks). That is, on average, 91% of NTC individuals were correctly not predicted to have an ASD diagnosis, and 16% of ASD individuals were correctly predicted to have an ASD diagnosis. ***Figure 1*** shows an overview of the bootstrap PPV across networks. We thus showed that high confidence predictions of ASD diagnosis made by individual brain networks did not lead to predictions with high PPV.

### Functionally similar brain networks predict correlated conformal scores

We investigated whether groups of brain networks existed that gave rise to similar conformal predictions of ASD diagnosis and could be combined to achieve more accurate group predictions. To do so, we computed the correlation between the ASD conformal scores predicted by the individual brain network predictors and applied hierarchical agglomerative clustering to derive 7 groups of networks with correlated conformal scores:

- group 1 was a single network group of the fronto-parietal network
- group 2 combined limbic and temporal networks (orbito-frontal cortex, inferior temporal sulcus, lateral default mode network, and amygdala-hippocampal complex)
- group 3 was a single network group containing the basal ganglia network
- group 4 combined sub-components of the default mode network (anterior-, and posteriormedial default mode network, and perigenual anterior cingulate and ventromedial prefrontal cortex)
- group 5 combined unimodal sensory networks (ventral, and dorsal somatomotor network, and auditory network)
- group 6 combined attention networks (medial ventral, and lateral ventral attention network, and fronto parietal task control network)
- group 7 combined visual networks (medial-, lateral-, and downstream visual network).

We thus showed that functionally similar brain networks tended to give rise to correlated conformal predictions of ASD diagnosis.

We combined the conformal scores predicted by brain networks within each group to generate high confidence group predictions of ASD diagnosis and evaluated them over 100 random bootstrap samples (see Methods for detailed explanation of the process of combining conformal scores and the corresponding adjustment of the conformal score thresholds). The average bootstrap PPV across all groups of networks was 64% with high specificity (90%) and low sensitivity (17%). The bootstrap PPV of groups of networks was generally close to that of the average bootstrap accuracy across the individual networks within them:

- *PPV*_*group*4_ = 69.7% compared to an average 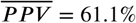 of the individual networks
- *PPV*_*group*4_ = 69.2% compared to 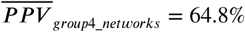
- *PPV*_*group*5_ = 64.8% vs 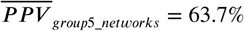
- *PPV*_*group*6_ = 59.0% vs *PPV*_*group*6_*networks*_ = 65.4%
- *PPV*_*group*7_ = 63.4% vs 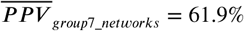

Conformal scores for the two single network groups (group 1 and 3) were adjusted identically to those of the multi-network groups, which resulted in altered bootstrap PPV estimates:

- *PPV*_*group*1_ = 61.4% vs *PPV*_*FP*_*network*_ = 63.8%
- *PPV*_*group*3_ = 60.6% vs *PPV*_*basal*_*ganglia*_ = 63.9%

We thus show that groups of brain networks with correlated conformal scores predicted ASD diagnosis with only marginally higher PPV than individual brain networks.

### Ensemble of transmodal networks forms high risk ASD signature

We further combined brain networks with correlated conformal scores into two large ensemble predictors and investigated whether they gave rise to distinct high risk signatures of ASD diagnosis. The first ensemble combined conformal scores of the nine more transmodal brain networks from groups 1 (fronto-parietal), 2 (limbic), 3 (basal ganglia), and 4 (default mode network). The second ensemble combined conformal scores of the remaining nine more unimodal brain networks from group 5 (sensorimotor), group 6 (attention), and group 7 (visual). We evaluated the predictions of these two ensemble models across 100 random bootstrap samples. The bootstrap PPV of the combined conformal scores in ensemble 1 was 88.7%, considerably higher than the average of the corresponding group predictors (62.9%). The bootstrap PPV in ensemble 2 was 72.0%, compared to the average PPV of the corresponding group predictors of 64.6%. Ensemble 1 predicted ASD diagnosis with higher specificity (99.5%) and lower sensitivity (4.9%) than ensemble 2 (specificity 97.1%, sensitivity 7.4%). Combining all brain networks into a whole brain model did not improve the PPV (average bootstrap PPV 76.6%). Based on these findings, we chose to further investigate the high PPV signature of ensemble 1 in the independent replication data set. We thus show that combining correlated conformal predictions of individual brain networks into ensemble predictors gave rise to a single FC based high risk signature (HRS) of ASD diagnosis.

### High risk ASD signature generalizes to independent data

We explored the performance of the HRS of ensemble 1 in an independent replication sample to determine its generalizability. For each individual in the replication sample, we thus computed the conformal scores for the ASD and NTC label with respect to the individuals in the discovery sample. The HRS identified 10 individuals from 6 different imaging sites (USM_1: 3, GU_1: 3, NYU_1: 1, SDSU_1: 1, IP_1: 1, KKI_1: 1) in the replication sample, and of those, 9 did have an ASD diagnosis. The PPV of the HRS of ensemble 1 was thus 90% in the replication sample, close to the average bootstrap PPV of 88.7% in the discovery sample. Specificity and sensitivity of the predictions were also similar to those estimated in the discovery sample: specificity = 99.5% (discovery: 99.5%), sensitivity = 4.2% (discovery: 4.9%). Predictions of the ensemble 2 model in the replication sample likewise performed similarly to bootstrap estimates in the discovery sample: PPV = 62.5% (discovery: 72.0%), specificity = 95.8% (discovery: 97.1%), sensitivity = 7.1% (discovery: 7.4%). We thus show that the high risk ASD signature identified in the discovery dataset generalized to an independent validation dataset with similar predictive performance.

### High risk ASD signature translates to 7-fold risk increase in general population

The discovery and replication samples were balanced to have equal numbers of individuals with ASD and NTC labels (i.e. the prevalence of ASD in our samples was 50%) in order to facilitate training and evaluation of the predictive models. However, the prevalence of ASD in an unselected population is estimated to be much lower (i.e. about 1 individual with ASD for 89 NTCs). The HRS correctly identified 4.2% of individuals with ASD in the sample (sensitivity) and incorrectly identified 0.5% of individuals with NTC (1 - specificity or false positive rate) in the sample. If the number of individuals with NTC considerably exceeds the number of individuals with ASD, the rate of individuals correctly identified by the model (accuracy), therefore also changes. To estimate the performance of the high risk signature in an unselected population, we thus computed the expected accuracy for a prevalence of ASD of 1/90 or 1.11%. In this context, the HRS correctly identified 0.046% of the population (4.2% sensitivity 1.11% individuals with ASD) and incorrectly identified 0.49% of the population (0.5% false positive rate ∗ 98.89% individuals without ASD or with NTC). Therefore the ratio of correctly identified individuals to all identified individuals (i.e. the PPV) was 0.046%/(0.046% + 0.49%) or 8.545%. An individual in the general population identified by the HRS thus had an estimated risk of ASD of 8.5% or a 7.7 fold increase in risk over the baseline risk in this population. We thus show that the high risk ASD signature conferred an estimated 7.7 fold increase in individual risk over the baseline.

### High risk ASD signature identifies individuals with severe symptoms

We next investigated the symptom characteristics of the individuals who were identified by the HRS model. To that end, we reported their ADOS severity measures and compared them to those of unselected individuals from the same clinical category. Because only 10 individuals were identified by the HRS model, these results are exploratory and we limited ourselves to reporting only descriptive measures. Calibrated ADOS severity scores (ADOS-CSS) would have been the preferred measure to interpret symptom severity because of their standardized range (from 1: least severe symptoms to 10: most severe symptoms), and because of their comparability across ADOS modules and across different ages. However, ADOS-CSS were only available for 3 identified individuals. Using a previously published technique we therefore computed proxy ADOS-CSS based on the available data (see Methods for details). We reported these closely approximated (r = 0.94) proxy ADOS-CSS together with the ADOS raw total scores.

The median of proxy ADOS-CSS was higher among the nine identified individuals with ASD (median = 9, interquartile range = 4 — 9) than among the remaining individuals with ASD who were not identified by the HRS model (median = 6, interquartile range = 5 — 8). The single NTC individual identified by the model had a higher proxy ADOS-CSS of 3 than the remaining NTC individuals who were not identified by the HRS model (median = 1, interquartile range = 1 — 1). The same comparison using raw ADOS total scores revealed an analogous finding: the median of raw ADOS total scores was higher among the nine identified ASD individuals (median = 15, interquartile range = 13 — 17) than among the remaining unidentified ASD individuals (median = 10, interquartile range = 8 — 13.25). Accordingly, the single identified NTC individual had a higher raw ADOS total score of 7 than the remaining unidentified NTC individuals (median = 1, interquartile range = 0 — 2). ***Figure 3*** shows both the proxy ADOS-CSS and the raw ADOS total scores of the identified individuals compared to those of unidentified individuals with the same diagnostic class. Our exploratory findings thus indicate that the identified individuals showed particularly severe symptoms for their diagnostic class.

**Figure 3.**
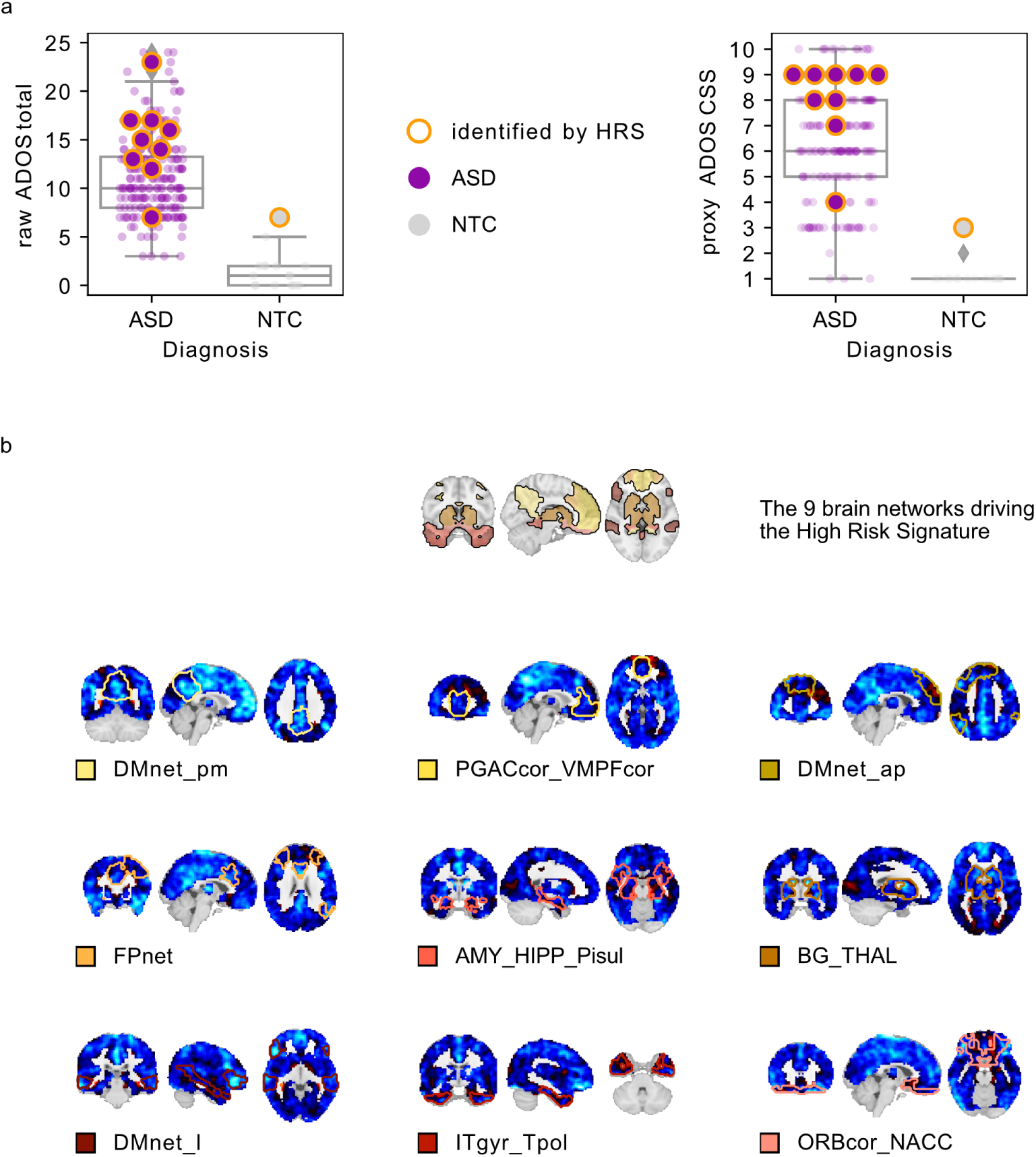
The HRS identifies individuals with severe symptoms and pervasive underconnectivity. a) Individuals identified by the high risk signature (circles with orange outline) have high proxy calibrated ADOS severity scores (left plot) and high raw ADOS total scores (right plot) compared to the average of their respective diagnostic category. Box-plots are based on individuals not identified by the HRS. b) The identified individuals share a pattern of distributed below average functional connectivity of the nine networks driving the high risk signature (the networks are denoted by name and coloured outline on their respective connectivity maps).

### High risk signature characterized by underconnectivity

To identify the FC pattern of the individuals detected by the HRS model, we investigated the average residual connectivity maps of the identified individuals for the nine brain networks contributing to the HRS signature. ***Figure 3***b shows the average residual connectivity maps of the nine networks. The average residual FC maps from all nine brain networks are characterized by pervasive under-connectivity with respect to the rest of the discovery sample. We thus show that the FC signatures of individuals identified by the HRS model were characterized by wide-spread underconnectivity of the nine involved brain networks with respect to the sample average.

### Conformal prediction not driven by nuisance covariates

To ensure that the conformal scores used to make the high confidence prediction were not driven by known sources of nuisance variance, we computed the Pearson’s correlation coefficient of ASD conformal scores with age and head motion across bootstrap samples in the discovery sample. Our results show that for all individual network predictors, the 90 % confidence interval of correlation coefficients with age and head motion included zero (see ***Figure 4***), and the median correlation coefficients were close to zero (age: network average r = –0.01, range: –0.03 — 0.03; head motion: network average r = -0.0027, range: –0.022 — 0.02). The ASD conformal scores of the two ensemble predictors similarly show median correlation estimates close to zero with age (r_ens1_ = –0.01; r_ens2_ = 0.01) and head motion (r_ens1_ = 0.01; r_ens2_ = –0.005) and the 90% confidence intervals of either correlation included zero (see ***Figure 4***). We thus conclude that the estimated ASD conformal scores were not driven substantially by nuisance covariates.

**Figure 4.**
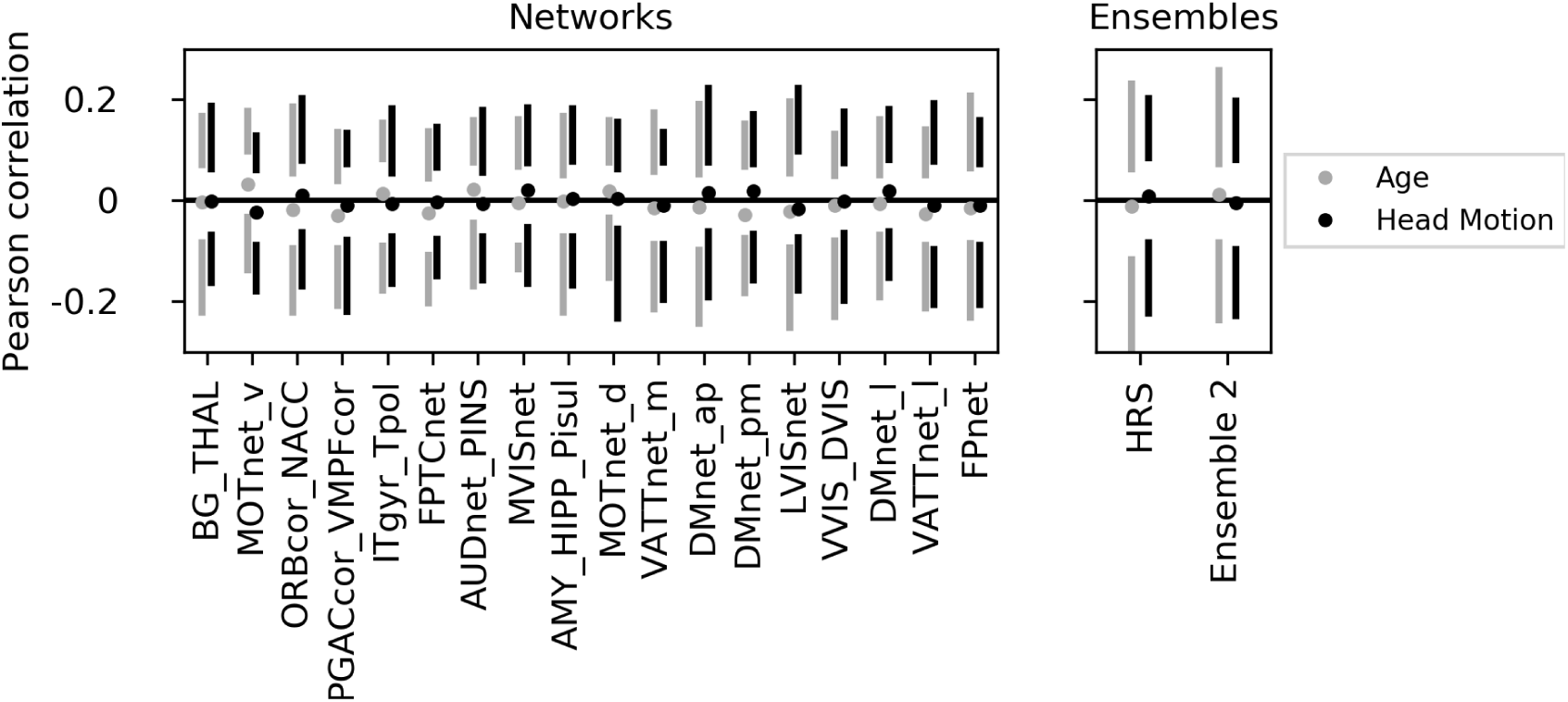
The conformal predictions are not driven by nuisance covariates. a) The distribution of correlations of ASD conformal scores predicted by individual networks (left) and the two ensemble models (right) with head motion (black) and age (grey) are shown across 100 bootstrap samples. Circles represent the median correlation score across bootstrap samples, vertical lines span the 5th to 25th percentile (lower bar) and 75th to 95th percentile (upper bar) of correlation scores respectively. All median correlation scores are close to zero and enclose zero within the 90% confidence interval.

### Conformal prediction performance exceeds baseline model

To determine if our FC based predictive signature performed better than a simple baseline model, we repeated the conformal prediction procedure using an individual’s age and in scanner head motion as input features. Following the same procedure described above, we then use the transductive conformal prediction approach to predict an ASD diagnosis only for those individuals in whom the model had high confidence. Our results show that such a baseline model did not predict ASD diagnosis with high confidence for any individuals in 90% of bootstrap samples (i.e. the sensitivity and PPV is zero). Among the 10% of bootstrap samples where the baseline model did make predictions, they were of high specificity (median = 100%) and low sensitivity (median = 7.9%) but low PPV (median = 50.5%, see ***Figure Supplement 1***). We thus show that the FC based network predictors performed better than a simple baseline model.

## Discussion

This work aimed to identify imaging risk endophenotypes of ASD that are both commonly found in the general population and confer a high risk of the disorder. We used a transductive conformal prediction approach to identify only those individuals for whom the ASD diagnosis could be predicted with high confidence on the basis of functional connectivity (FC). Our results showed that the combined predictions of nine brain networks gave rise to a single high risk FC-signature that identified individuals with severe symptoms and pervasive underconnectivity in an independent validation dataset. This FC-signature confers a more than 7-fold increased risk of ASD diagnosis in the general population where it is identified in an estimated 1 in 200 individuals, compared to a baseline ASD prevalence of 1 in 90 individuals. The risk conferred by our FC-signature constitutes a more than 3 and a half fold increase over current neuroimaging prediction models of ASD.

### Model performance

The multi-network risk FC-signature we have identified here confers a positive predictive value (PPV) of 8.5%, more than 7 times higher than the baseline risk of ASD diagnosis in the general population (1 in 90 ≈1.11%). This PPV is also more than a 3.5 fold larger than previously published imaging based prediction models for ASD. We achieved this considerable increase in individual risk by changing the goal of our prediction model. Whereas previous models have made predictions for all individuals in heterogeneous case-control populations, we limited predictions to only a subset of individuals for whom our model has very high confidence in an ASD diagnosis. Although our model made only few predictions, those predictions carry a much higher risk of an ASD diagnosis for the identified individuals. The result is a prediction with a much higher specificity (99.5% compared to 72.3% and 63% for traditional approaches, ***Heinsfeld et al., 2018***; ***Abraham et al., 2017***) and much lower sensitivity (4.2%, compared to 61% and 74% respectively). It is thus important to point out that here we have not proposed a better prediction learning model, but rather addressed a different objective. It is reasonable to assume that the conformal prediction approach would lead to predictions with similarly high specificity when applied to previously published imaging models.

In the general population, our high risk signature is estimated to be identified in about 1 in 200 individuals. It is thus approximately two orders of magnitude less common than ASD-related SNPs (***Grove et al., 2019***), that confer negligible individual risk, and about two orders of magnitude more common than rare monogenic syndromes (***de la Torre-Ubieta et al., 2016***), that confer very high risk of ASD (see ***Figure 2***). To the best of our knowledge, there are no other imaging or genetic risk signatures of autism that confer a comparable amount of individual risk and are still relatively common. Polygenic risk signatures of similar prevalence and risk have been identified recently for some common diseases (***Khera et al., 2018***) and may be identified in the future for ASD (***Martin et al., 2018***). However, the comparatively low number of identified common variants for ASD (i.e. only 5 ASD specific SNPs have been robustly identified to date, ***Grove et al., 2019***, compared to 108 that have been identified for schizophrenia, ***Schizophrenia Working Group of the Psychiatric Genomics Consortium 2014***) and the massive sample sizes required to robustly estimate polygenic risk (e.g. approximately two orders of magnitude larger than those used in this study) currently constitute important obstacles for these potential discoveries.

### The high risk signature is mainly driven by transmodal brain networks

Individually, the 18 brain networks did not predict ASD with high PPV. By clustering networks with correlated conformal scores and combining their predictions, we identified two equally sized sets of brain networks. The first one gave rise to the high risk ASD FC-signature and involved predominantly transmodal networks in the default mode and fronto parietal network, but also of subcortical areas (***Alves et al., 2019***). This is consistent with previous FC-based prediction models of ASD that found the most predictive functional connections to involve transmodal areas such as the temporal parietal junction and areas of the fronto-parietal control network (***Abraham et al., 2017***), connections within the cingulo-opercular network (***Yahata et al., 2016***) and of supramarginal, middle temporal, and cingulate gyri (***Heinsfeld et al., 2018***). FC alterations of transmodal networks have also been consistently reported in the ASD case-control literature (***Monk et al., 2009***; ***Holiga et al., 2019***; ***Just et al., 2007***), in particular in regions of the default mode network (***Washington et al., 2014***; ***Assaf et al., 2010***).

The second ensemble, that did not predict ASD with high PPV, predominantly consisted of unimodal networks in the visual, auditory, and somatosensory cortices involved in sensory processing, but also the ventral attention network. Although FC of primary sensory brain regions was previously found to be less predictive of ASD diagnosis than that of transmodal regions (***Heinsfeld et al., 2018***), there is extensive evidence of ASD related FC alterations in unimodal areas (***Oldehinkel et al., 2019***). Why then do we observe this difference in predictive performance between the two ensembles?

The distinction between the FC of unimodal and transmodal networks is a very robust and well established finding (***Raichle et al., 2001***; ***Fox et al., 2005***; ***Buckner and DiNicola, 2019***) that is also reflected in their opposing FC alterations in ASD. Whereas transmodal regions were found to be reproducibly over-connected in ASD, unimodal regions were found to be reproducibly underconnected in a recent multi-center study (***Holiga et al., 2019***). Transmodal and unimodal brain networks were recently shown to lie on opposite ends of a cortical gradient of functional hierarchy (***Margulies et al., 2016***) that is altered in ASD (***Hong et al., 2019***), suggesting a dysfunctional separation between primary sensory networks and the default mode network. It is thus possible that both ensembles capture distinct ASD risk signatures but only one of them could be reliably identified in our dataset.

### Individuals identified by the HRS have severe symptoms and functional underconnectivity

The high risk FC-signature identified a group of ten individuals from the independent validation dataset, and nine of them had a diagnosis of ASD. These individuals tended to also have high symptom severity measures. Notably, the one individual identified by the high risk signature who did not have an ASD diagnosis did also have unusually severe symptoms compared to other NTC individuals. This individual may reflect a broader autism phenotype that extends into the general population (***Baron-Cohen et al., 2001***) and is picked up by our model. It is possible that the high risk FC-signature identifies a subtype of ASD patients with particularly severe symptoms. Because these individuals are identified due to their strong dissimilarity with NTC, this interpretation would be consistent with a view of neurodevelopmental disorders as an extreme deviation from normal functioning (***Marquand et al., 2019***).

The identified individuals shared a profile of pervasive functional underconnectivity among the transmodal networks that gave rise to the high risk FC-signature. Although dysconnectivity of transmodal brain networks, and the default mode network in particular (***Monk et al., 2009***), have been consistently reported in the ASD case-control literature, the direction of these effects has not been consistent (***Padmanabhan et al., 2017***; ***Hull et al., 2016***) and both over- and under-connectivity have been related to increases in symptom severity (***Assaf et al., 2010***; ***Supekar et al., 2013***). Notably, the profile of transmodal network underconnectivity we have identified here stands in contrast to recent case-control findings of reproducible, ASD-related prefrontal and parietal overconnectivity in a large, multi-center study (***Holiga et al., 2019***). These contrasting findings may reflect the inherent limitations of case-control studies to identify subtypes of FC alterations that are strongly linked to ASD. Indeed, recent work on ASD related FC subtypes similarly found a profile of transmodal underconnectivity (***Tang et al., 2019***). Our results are compatible with reports by other imaging based prediction models of ASD that found underconnectivity between default mode network subregions to be the most discriminant features for prediction (***Abraham et al., 2017***; ***Heinsfeld et al., 2018***; ***Yahata et al., 2016***).

### Limitations

These findings have to be interpreted in light of their limitations: Our analyses only included male individuals which is a common problem in the field (***Khundrakpam et al., 2017***; ***Hong et al., 2019***) due to the higher frequency with which ASD is diagnosed among male individuals (***Lai et al., 2014***). Recent data curation efforts have therefore started to deliberately include more female individuals (***Di Martino et al., 2017***; ***Bedford et al., 2019***).

The behavioural and symptomatic characterization of individuals detected by the high risk signature were limited by the inconsistent availability of phenotypic information in our data. A future comprehensive characterization of the high risk signature will have to make use of large scale datasets with more complete phenotyping and will help better clarify the neurobiologically defined subset of at risk individuals in terms of their cognitive and symptom profile.

Due to the transductive nature of our conformal prediction model, we can only control for nuisance covariates that are available both in the reference sample and for the predicted individual. We could therefore not regress effects due to recording site from the individual FC data. Despite this fact, the high risk ASD signature identified individuals from across different imaging sites with high PPV in an independent dataset, suggesting that the identified FC endophenotype is robust to site differences (see also ***Orban et al., 2018***).

We have estimated the general population risk conferred by our high risk signature based on its performance in the independent dataset. In line with our expectations, only very few individuals were identified. Risk signatures with such a low prevalence are typically validated on much larger datasets to ensure the robustness of the performance estimates (***Khera et al., 2018***). General population samples of this magnitude that also provide imaging data have recently become available (***Bycroft et al., 2018***) and validating the high risk signature on these data is a natural next step to establish robust estimates of the prediction performance of this high risk signature.

### Future directions

The high risk FC-signature that we have described here provides interesting implications for future research. As a cohort of individuals with similar FC alterations at high risk of an ASD diagnosis, our signature identifies a potential population of interest to investigate the link between neurobiological aberration, behavioural symptoms and genetic mechanisms. It may thus provide a starting point to disentangle the heterogeneous relationships across these levels of research in ASD (***Lombardo et al., 2019***). An important next step will be to investigate the stability of this FC signature across time (***Jacob et al., 2019***) and to establish at what point of the developmental trajectory it can be differentiated (***Emerson et al., 2017***). These questions will require large scale longitudinal data of at risk individuals, such as the IBIS dataset (***Wolff et al., 2012***). Finally, investigating this high risk ASD signature in other, comorbid (***Simonoff et al., 2008***) neurodevelopmental disorders may help clarify the symptomatic (***Grzadzinski et al., 2011***), neurobiological (***van den Heuvel and Sporns, 2019***; ***de Lange et al., 2019***), and genetic (***Cross-Disorder Group of the Psychiatric Genomics Consortium et al., 2013***) overlap between these disorders and the autism spectrum.

### Conclusion

We have identified a functional connectivity endophenotype associated with high risk of ASD that can be detected with high positive predictive value in independent data. Decomposing the autism spectrum bit by bit in this manner may eventually help us understand the multitude of etiological pathways and their extension to the general population.

## Methods and Materials

### Sample

All data were sampled from the ABIDE 1 (***Di Martino et al., 2014***) and ABIDE 2 (***Di Martino et al., 2017***) dataset releases that contain imaging data for ASD patients and NTC. We used the ABIDE 1 release as a discovery dataset and retained the ABIDE 2 release as an independent validation dataset.

The final discovery dataset consisted of 478 male individuals (*Age* = 16.67 (6.67), *N*_*ASD*_ = 239) from 13 recording sites. From the complete ABIDE 1 dataset of 1112 individuals (*Age* = 17.04 (8.04), *N*_*ASD*_ = 539) from 20 imaging sites we excluded 164 female individuals due to strong sex imbalance. Of the remaining sample, 557 individuals from 13 imaging sites were successfully preprocessed and passed visual quality control (*Age* = 16.65 (6.75), *N*_*ASD*_ = 272). In order to control for the effects of nuisance covariates in the data without removing variance due to the ASD diagnosis, we then matched ASD and NTC individuals on age and head motion within each imaging site by propensity score matching without replacement (***Rosenbaum and Rubin, 1985***).

The validation dataset consisted of 424 male individuals (*Age* = 13.66 (5.25), *N*_*ASD*_ = 212) from 16 imaging sites. From the complete ABIDE 2 dataset of 1114 individuals (*Age* = 14.86 (9.16), *N*_*ASD*_ = 521) from 19 imaging sites, we excluded 258 female individuals due to the strong sex imbalance and to match the sample characteristics of the discovery sample. Of the remaining sample, 587 (*Age* = 13.94 (5.9), *N*_*ASD*_ = 273) from 16 imaging sites were successfully preprocessed and passed visual quality control. In line with the sample selection of the discovery sample, we then matched ASD and NTC individuals on age and head motion within each imaging site using propensity score matching without replacement.

### Clinical diagnosis and severity estimates

The individuals from the ABIDE 1 and ABIDE 2 samples included in this study were diagnosed with ASD by expert clinicians based on either the Autism Diagnostic Observation Schedule (ADOS) (***Lord et al., 2000***; ***Gotham et al., 2007***; ***Lord et al., 2012***) or the Autism Diagnostic Interview - Revised (***Lord et al., 1994***). ADOS total scores are available for 228 (*N*_*ASD*_ = 196) individuals in the discovery sample and 226 (*N*_*ASD*_ = 209) individuals in the validation sample. Although higher ADOS total scores indicate more serious impairments, ADOS raw total scores were not originally intended to compare individuals with different ages, or tested with different ADOS modules. For this purpose, the original authors provide a standardized method (***Gotham et al., 2009***) to convert ADOS total scores to 10-point calibrated severity scores (10 being the most severe), which are less influenced by an individuals’ age and other demographic confounds. However, ADOS-CSS were not available for many individuals in the discovery (*N* = 107, *N*_*ASD*_ = 91) and validation sample (*N*_*ASD*_ = 115). In order to better contextualize the symptom severity of individuals from different age groups, we computed proxy ADOS-CSS scores by using the available ADOS total scores and the published conversion table (***Moradi et al., 2017***). Proxy ADOS-CSS scores were strongly correlated with true ADOS-CSS scores in both the discovery (*r* = 0.90) and the validation (*r* = 0.94) sample. Proxy ADOS-CSS scores could be computed for 221 individuals (*N*_*ASD*_ = 190) in the discovery and 223 (*N*_*ASD*_ = 207) in the validation sample.

### Imaging data preprocessing

Imaging data from individuals in both the discovery and independent validation sample underwent identical preprocessing through the NeuroImaging Analysis Kit (NIAK) version 1.13 (***Bellec et al., 2011***) running inside a Singularity (version 2.6.1) containerized environment (***Kurtzer et al., 2017***) and using an established in-house processing pipeline. In short, functional time series were corrected for in-scanner head motion and registered to the MNI152 stereotaxic space (***Evans et al., 1994***). Slow time drift signals were modeled on the continuous time series by a discrete cosine transformation and removed after censoring of time frames with excessive (> 0.4mm) head motion (***Power et al., 2012***), together with nuisance covariates of the average white matter, and cerebrospinal fluid signals, and the first principal components (accounting for 95% of variance) of the six degrees of freedom head motion estimates and their squares (***Giove et al., 2009***).

### Imaging data quality control

The preprocessed imaging data were visually quality controlled (QCed) to ensure the quality of the data. The QC was performed by a trained rater according to our in-lab standardized QC protocol (***Benhajali et al., 2019***) using a guided QC environment (***Urchs et al., 2018***). Imaging data were excluded from subsequent analyses in cases of failed brain extraction or coregistration to the stereotaxic space, visible motion artifacts, incomplete brain coverage of the field of view, or if fewer than 50 time frames remained after motion censoring. A large number of individuals from both the discovery and the validation dataset were found to have incomplete coverage of the cerebellum. In order to retain these otherwise correctly preprocessed individuals, we decided to exclude the cerebellum from the FC analyses.

### Functional connectivity estimation

Seed to voxel FC was estimated for functional brain networks defined in the MIST_20 atlas (***Urchs et al., 2017***). The MIST_20 atlas represents large, spatially distributed subcomponents of canonical FC networks. Of the 20 brain networks defined in the MIST_20 atlas, 2 were part of the cerebellum and were excluded (see above). For each of the remaining 18 brain networks, the average within-network time series was correlated with the time series of all non-cerebellar voxels using Pearson’s correlation. The FC organization of every individual in the discovery and validation was thus described by 18 network to voxel maps.

### High confidence classification

The transductive conformal classification (TCC) approach (***Vovk et al., 2005***; ***Nouretdinov et al., 2011***), which we have applied here, calculates the degree to which a new datapoint “conforms” to already classified data points in some measure of interest. The already classified data points were the reference set of ASD and NTC individuals of the discovery dataset, and our measure of interest was the FC of the 18 brain networks.

In contrast to an inductive classification approach, where a statistical model is first learned based on the properties of the reference set and then applied to new data, in a transductive classification, no model is learned and each new individual is classified directly and separately by comparing it to the properties of each class (ASD and NTC) in the reference set, and choosing the class (or classes) it most conforms to; see Chapelle, et al. (***Chapelle et al., 2006***) for a treatment regarding the difference between inductive and transductive learning. Each unclassified individual therefore has to be treated in the exact same way to ensure the independence of each classification.

#### Regression of nuisance covariates

We combined the unclassified individual and the reference sample and removed the group level average connectivity and the linear effect of age and head motion from the network FC maps.

#### Dimensionality reduction

Previous works have shown the capacity of FC subtypes to capture disease-related FC variability; e.g. (***Easson et al., 2019***). We therefore identified the 5 subtypes of FC variability across both the unclassified individual and the reference sample by hierarchical agglomerative clustering of spatially correlated, individual FC maps. For each individual we then computed the spatial similarity with the average FC map of each of the 5 FC subtypes.

#### Estimation of conformality and classification

The individual conformality estimate for either clinical label (i.e., ASD or NTC) was then computed similarly to the previous work of Nouretdinov et al. (***Nouretdinov et al., 2011***). In short, we first assumed an ASD label for each unclassified individual and then fit a logistic regression to predict ASD for both the unclassified individual and the reference sample, using the previously estimated similarity with FC subtypes as features. To reflect the fact that we wanted the model to make as few false positive errors as possible, we weighted the predicted values of ASD individuals by a large scaling factor (*w*_*ASD*_ = 10^16^). This was in line with the suggestion from the discussion of ***Nouretdinov et al.*** (***2011***). This forced the prediction model to only be concerned with the identification of ASD cases, with high specificity, at the expense of possible identification of NTC individuals. We computed the ASD conformal score for each unclassified individual as the percentage of ASD individuals in the reference sample with a predicted value equal to or smaller than the one that was predicted for that unclassified individual. In other words: if most ASD individuals had larger predicted values than the unclassified individual, then the unclassified individual did not conform to the ASD cohort and was an unusual ASD case, and thus the ASD conformal score would have been small due to the individual not “conforming” to the reference cohort of ASD individuals. An analogous process was then repeated to compute the NTC conformal score of the unclassified individual.

We rejected a clinical label (i.e. ASD or NTC) if the corresponding estimated conformal score was below a critical threshold of 5%. We predicted ASD with high confidence for only those individuals who had NTC conformal scores below the critical threshold and ASD conformal scores equal or greater than the critical threshold.

### Assessment of prediction performance

To assess the quality of the classification, we computed the sensitivity and specificity across the predicted individuals. The sensitivity of the classification:

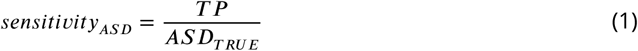

reflects the ability of our model to correctly predict ASD (TP) among those individuals who truly have ASD (ASD_TRUE_). Incorrectly predicting an ASD diagnosis for an individual without the diagnosis is known as a false positive (FP) error. Our approach tried to minimize the false positive error. The specificity of the classification:

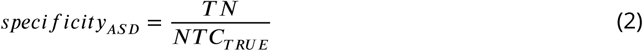

likewise reflects its ability to correctly not predict ASD (TN) among those individuals who truly do not have an ASD diagnosis (NTC_TRUE_). Incorrectly predicting an ASD individual as “not ASD” is known as a false negative (FN) error. The positive predictive value (PPV):

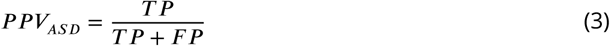

is the ratio of correct ASD predictions (TP) among all ASD predictions made by our model. It thus reflects the risk of an individual classified as ASD by our model to truly have an ASD diagnosis. Our approach aimed to maximize the positive predictive value. The PPV depends on the ratio of ASD_TRUE_ among all individuals in our sample. This ratio is known as the prevalence of ASD in the sample.

For an individual who was identified by the model as suspected ASD, the PPV_ASD_ provides an estimate of the individual probability of a true ASD diagnosis. If the model confers any risk, then the risk of ASD is larger for someone identified by the model than for someone not identified by the model. This measure is called the risk ratio (RR_ASD_):

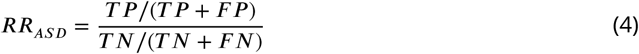

A similar metric that is independent of the prevalence of the disorder is the Odds ratio (OR). The odds of a true ASD diagnosis for a selected individual is the ratio of the probability

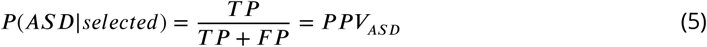

over the probability

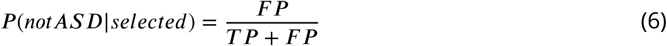

Both can be simplified to

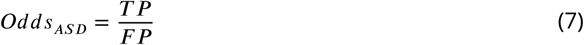

Analogous to the risk ratio, the Odds ratio (OR):

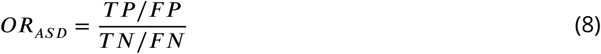

then reflects the ratio of odds of an ASD diagnosis for selected individuals over the odds of ASD for unselected individuals.

For a model that conveys no information on the ASD diagnosis, the odds of a true ASD diagnosis are the same for individuals who are identified by the model and for those who are not identified (i.e. the OR is 1).

An ideal model would correctly classify all individuals with ASD. That is, the set of selected individuals and individuals with ASD would be exactly overlapping. In practice, models with high PPV (e.g. monogenic risk markers) tend to select only a very small subset of individuals (low sensitivity) and models with high sensitivity tend to incorrectly select many individuals without ASD (low specificity, see ***Figure 5***). We can thus use the overlap between individuals with ASD and selected individuals to determine how close the model is to an optimal tradeoff between sensitivity and specificity. The Sørensen–Dice coefficient:

**Figure 5.**
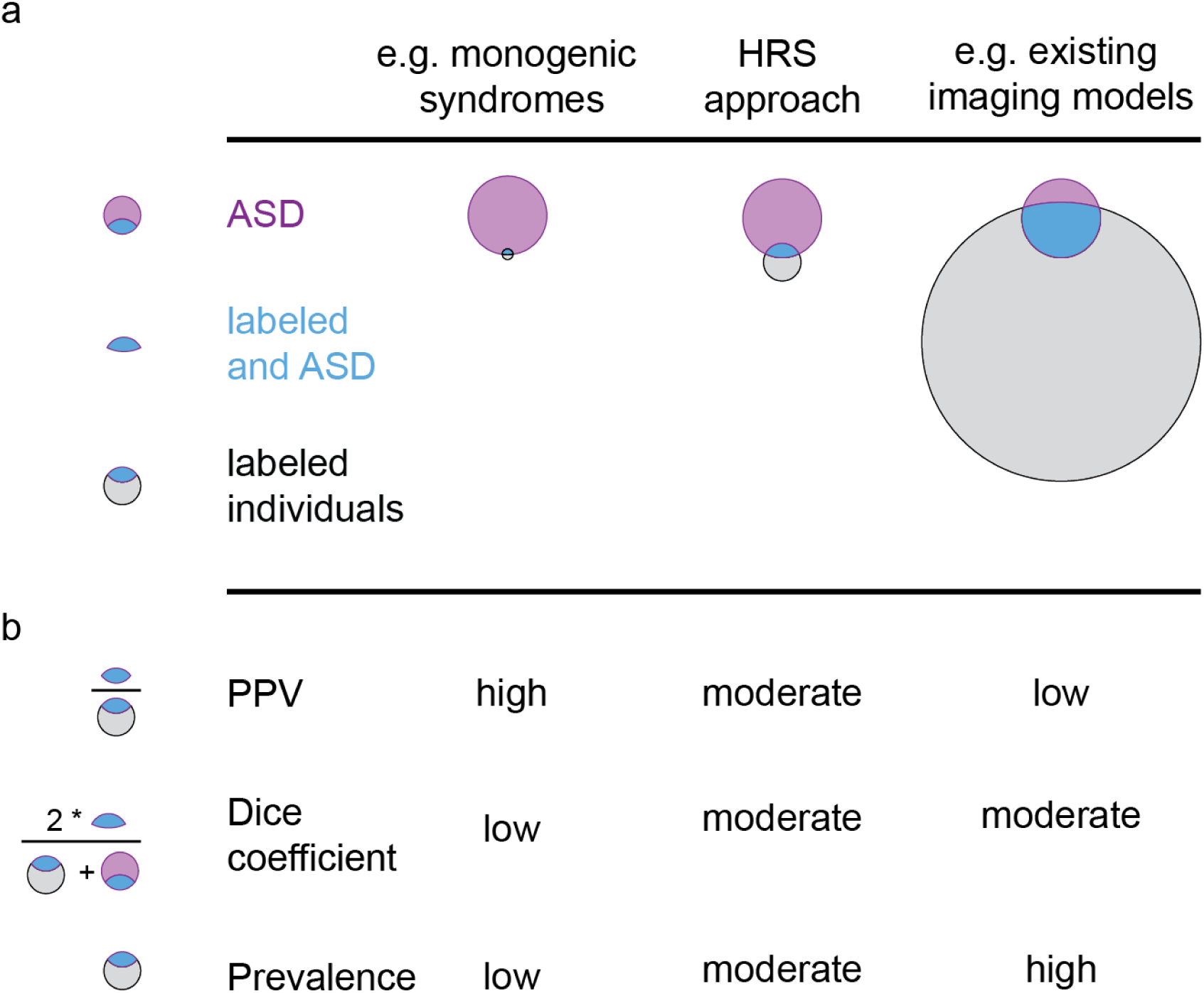
Schematic representation of properties of different ASD risk markers. a) A set of individuals in the population is found to express the risk marker (grey circle) and is thus labeled. Among the set of individuals with ASD in the population (purple circle), some are also labeled by the risk marker (blue region). Risk markers differ in the amount of labeled individuals from very few (left column) to very many (right column). b) Different metrics exist to evaluate the performance of the risk marker. The ratio of ASD individuals among the labeled individuals (PPV, see ***Equation 3***) can be very high if only a very few individuals are labeled by the risk marker (e.g. in monogenic syndromes with high risk for ASD, left column). However, the degree of congruence of ASD and labeled individuals (Dice, see ***Equation 9***) would be very low, because of the large number of unlabeled ASD individuals. Conversely, a risk marker that labels very many individuals may capture more ASD individuals and have a moderately higher Dice coefficient, but would have a very low ratio of ASD to labeled individuals (PPV) and thus confer very low individual risk (e.g. existing imaging based models, right column). The HRS approach presented here labels fewer individuals than current imaging models but those individuals are more likely to have ASD, resulting in higher PPV and comparable Dice coefficients.

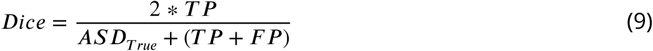

measures the ratio of correctly selected individuals over the sum of individuals with ASD (*ASD*_*True*_) and all selected individuals. It thus ranges between 0 (if the two sets are not overlapping) and 1 (if the two sets are completely overlapping).

### Bootstrap estimation

We estimated the model performance of each brain network predictor through bootstrap subsampling of the discovery data set. We drew two random bootstrap samples from the discovery data set and assigned one to be the reference data set and the other to be the prediction data set. The ASD diagnosis of each individual in the prediction data set was then separately predicted based on the individuals in the reference data set, following preprocessing, feature extraction and training as described above. We repeated this process 100 times for each brain network and computed the average performance metrics of each predictor across bootstraps (see, e.g. ***Efron, 1983***, regarding bootstrap predictor evaluation methods).

### Combination of correlated conformal predictions

To identify similarities of conformal predictions between the 18 functional brain networks, we computed the pairwise correlation of ASD non-conformity on the discovery sample. We then used hierarchical agglomerative clustering to identify groups of networks with correlated ASD conformal score estimates. We selected a 7 and 2 cluster solution based on a visual inspection of the network by network correlation matrix. Within each cluster of networks, conformal score estimates (i.e. probability estimates of non-conformity with each class label) were combined using the p-value averaging methods of (***Vovk and Wang, 2012***). Specifically, we averaged over the p-values that are associated within each network using the squared-mean merging function, which produces a valid aggregate p-value from the combination of any finite number of potentially correlated individual p-values. This requirement of validity is important in order to maintain the conformity properties when using these cluster-aggregated p-values as inputs in a conformal predictor.

The aggregation of p-values in the discovery sample was observed to average over the information that are inherent in each of the contributing p-values. As such, less informative network elements tended to decrease the explanatory power of the more informative elements. The overall effect was that the cluster non-conformity threshold tended to be conservative in identifying interesting observations, when compared to the same threshold value, applied to individual networks. In order to mitigate against this conservative effect, we used a more liberal threshold for cluster-aggregated p-values, than those used for individual networks. That is, we adjusted the critical non-conformal threshold to 0.2 from 0.05.

### Validation on the independent dataset

The HRS identified on the discovery sample was then validated on the independent validation sample. To do so, the ASD and NTC non-conformity estimate of each individual in the validation sample was computed by using the individuals of the discovery sample as the reference cohort. Each individual in the validation sample was predicted independently after group level nuisance regression and dimensionality reduction with respect to the reference sample.

### Estimation of model performance in the general population

The discovery and validation sample had equal rates of ASD patients and NTC individuals (i.e. 1 ASD for each 1 NTC). The prevalence of ASD in the general population is however much lower (1 ASD for each 89 NTC). Based on the estimated specificity and sensitivity of our model in the independent validation sample, we estimated the positive predictive value (*PPV*_*ASD*_) of the HRS in the general population.

## Acknowledgments

This research was supported by computation resources of Calcul Quebec and Compute Canada. We thank Yu Zhang and Gleb Bezgin for helpful discussions. For their feedback on the writing of this manuscript we want to thank Julie Boyle, and Jonas Nitschke. We thank the ABIDE consortium for making publicly available the large datasets that this study was based on.

---

**Figure 4–Figure supplement 1.**
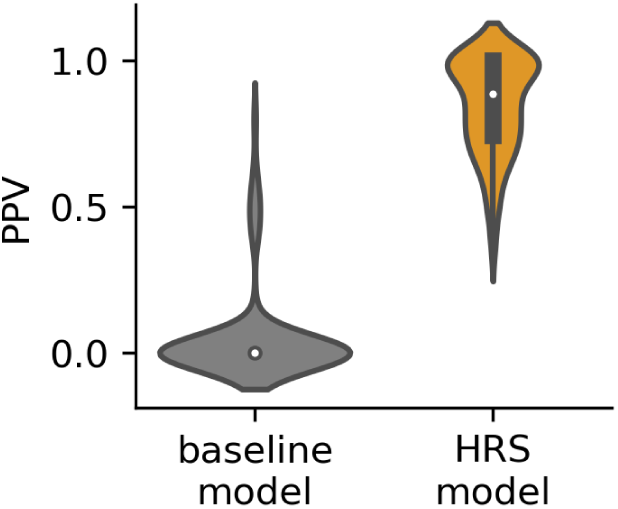
Predictions by the HRS exceed the PPV of those by a simple baseline model. The distribution of PPV estimates across 100 bootstrap samples is denoted by violin plots for each model

**Figure 4–Figure supplement 2.**
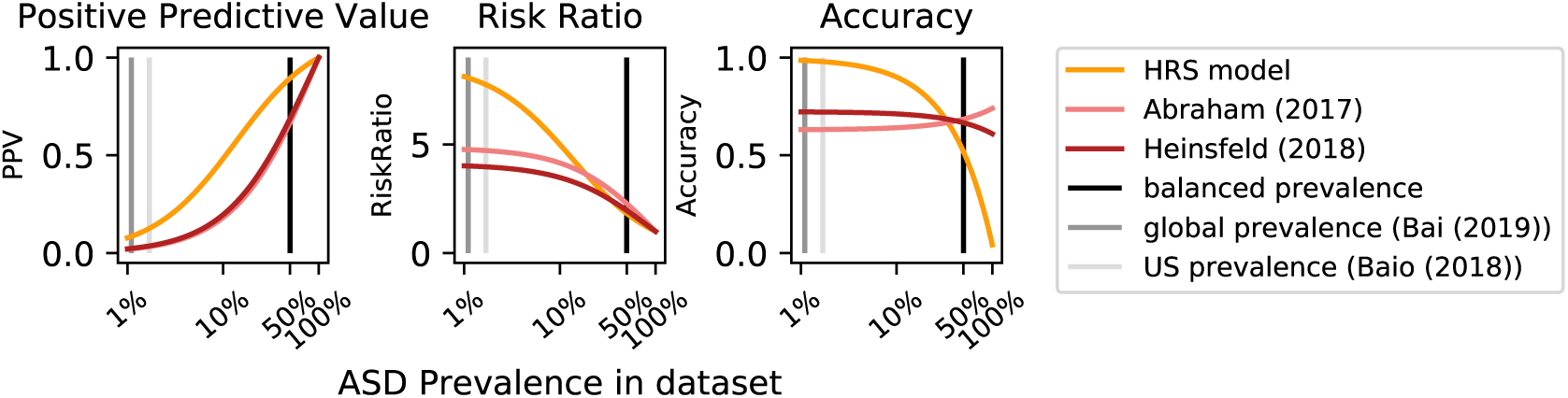
The impact of different levels of ASD prevalence in the data are shown for different metrics that are commonly used to evaluate prediction models. In balanced samples (black vertical line) that are commonly used to train models, traditional models (pink lines) that balance sensitivity and specificity achieve high accuracy. However, predictions by traditional models confer lower individual risk (PPV), particularly for low ASD prevalence, close to the baseline rate in the general population (grey lines)

